# Induction of Muscle Regenerative Multipotent Stem Cells from Human Adipocytes by PDGF-AB and 5-Azacytidine

**DOI:** 10.1101/2020.06.04.130872

**Authors:** Avani Yeola, Shruthi Subramanian, Rema A. Oliver, Christine A. Lucas, Julie A. I. Thoms, Feng Yan, Jake Olivier, Diego Chacon, Melinda L. Tursky, Tzongtyng Hung, Carl Power, Philip Hardy, David D. Ma, Joshua McCarroll, Maria Kavallaris, Luke B. Hesson, Dominik Beck, David J. Curtis, Jason W.H. Wong, Edna C. Hardeman, William R. Walsh, Ralph Mobbs, Vashe Chandrakanthan, John E. Pimanda

**Author notes:** corresponding authors Dr. Vashe Chandrakanthan, Dr. John Pimanda.

## Abstract

Terminally differentiated murine osteocytes and adipocytes can be reprogrammed using platelet-derived growth factor–AB and 5-Azacytidine into multipotent stem cells with stromal cell characteristics. To generate a product that is amenable for therapeutic application, we have modified and optimised culture conditions to reprogram human adipocytes into induced multipotent stem cells (iMS) and expand them *in vitro*. The basal transcriptomes of adipocyte-derived iMS cells and matched adipose-tissue-derived mesenchymal stem cells were remarkably similar. However, there were distinct changes in histone modifications and CpG methylation at *cis-*regulatory regions consistent with an epigenetic landscape that was primed for tissue development and differentiation. In a non-specific tissue injury xenograft model, iMS cells contributed directly to new muscle, bone, cartilage and blood vessels with no evidence of teratogenic potential. In a cardiotoxin muscle injury model, iMS cells contributed specifically to satellite cells and myofibres without ectopic tissue formation. Taken together, human adipocyte derived iMS cells regenerate tissues in a context dependent manner without ectopic or neoplastic growth.

## INTRODUCTION

The goal of regenerative medicine is to restore function by reconstituting dysfunctional tissues. Most tissues have a reservoir of tissue resident stem cells with restricted cell fates suited to the regeneration of the tissue in which they reside (*1-4*). The innate regenerative capacity of a tissue is broadly related to the basal rate of tissue turnover, the health of resident stem cells and the hostility of the local environment. Bone marrow transplants and tissue grafts are frequently used in clinical practice but for most tissues, harvesting and expanding stem and progenitor cells is currently not a viable option (*5, 6*). Given these constraints, research efforts have been focussed on converting terminally differentiated cells into pluripotent or lineage restricted stem cells (*7, 8*). However, tissues are often a complex mix of diverse cell types that are derived from distinct stem cells. Therefore, multipotent stem cells may have advantages over tissue specific stem cells. To be of use in regenerative medicine, such cells would need to respond appropriately to regional cues and participate in context dependent tissue regeneration without forming ectopic tissues or teratomas. Mesenchymal stem cells (MSCs) were thought to possesses some of these characteristics (*9-11*) but despite numerous on-going clinical trials, evidence for their direct contribution to new tissue formation in humans is sparse, either due to the lack of sufficient means to trace cell fate in hosts *in vivo* or failure of these cells to regenerate tissues (*12, 13*).

We previously reported a method by which primary terminally differentiated somatic cells could be converted into multipotent stem cells, which we termed induced multipotent stem cells (iMS) (*14*). These cells were generated by transiently culturing primary mouse osteocytes in medium supplemented with azacitidine (AZA; 2-days) and platelet derived growth factor-AB (PDGF-AB; 8-days). Although the precise mechanisms by which these agents promoted cell conversion was unclear, the net effect was reduced DNA methylation at the OCT4 promoter and re-expression of pluripotency factors (OCT4, KLF4, SOX2, c-MYC, SSEA-1 and NANOG) in 2-4% of treated osteocytes. iMS cells resembled MSCs with comparable morphology, cell surface phenotype, colony forming unit-fibroblast (CFU-F), long-term growth, clonogenicity and multi-lineage *in vitro* differentiation potential. Importantly, iMS cells also contributed directly to *in vivo* tissue regeneration and did so in a context dependent manner without forming teratomas. In proof of principle experiments, we also showed that primary mouse and human adipocytes could be converted into long-term repopulating CFU-Fs by this method using a suitably modified protocol (*14*).

AZA, one of the agents used in this protocol, is a cytidine nucleoside analogue and a DNA hypomethylating agent that is routinely used in clinical practice for patients with higher risk myelodysplastic syndrome (MDS) and for elderly patients with acute myeloid leukemia (AML) who are intolerant to intensive chemotherapy (*15, 16*). AZA is incorporated primarily into RNA, disrupting transcription and protein synthesis. However, 10-35% of drug is incorporated into DNA resulting in the entrapment and depletion of DNA methyltransferases (DNMT) and suppression of DNA methylation (*17*). Although the relationship between DNA hypomethylation and therapeutic efficacy in MDS/AML is unclear, AZA is known to induce an interferon response and apoptosis in proliferating cells (*18-20*). PDGF-AB, the other critical reprogramming agent is one of five PDGF isoforms (PDGF-AA, PDGF-AB, PDGF-BB, PDGF-CC and PDGF-DD), which bind to one of two PDGF receptors (PDGFRα and PDGFRβ) (*21*). PDGF isoforms are potent mitogens for mesenchymal cells and recombinant human PDGF-BB is used as an osteoinductive agent in the clinic (*22*). PDGF-AB binds preferentially to PDGFRα and induces PDGFR-αα homodimers or -αβ heterodimers. These are activated by autophosphorylation to create docking sites for a variety of downstream signaling molecules (*23*). Although we have previously demonstrated induction of CFU-Fs from human adipocytes using PDGF-AB/AZA (*14*), the molecular changes which underly conversion, and the multi-lineage differentiation potential and *in vivo* regenerative capacity of the converted cells have not been determined.

Here we report an optimised PDGF-AB/AZA treatment protocol that was used to convert primary human adipocytes from adult donors aged 27-66 years into iMS cells with long-term repopulating capacity and multi-lineage differentiation potential. We also report the molecular landscape of these human iMS cells along with that of MSCs derived from matched adipose tissues, and the comparative *in vivo* regenerative and teratogenic potential of these cells in mouse xenograft models.

## RESULTS

### Generating iMS cells from primary human adipocytes

Primary mature human adipocytes were harvested from subcutaneous fat (Figure 1A; Table S1) and their purity confirmed by flow cytometry with specific attention to the absence of contaminating adipose-derived mesenchymal stem cells (AdMSCs) (Figure S1A). As previously described (*14*), plastic adherent adipocytes were cultured in αMEM containing 200ng/mL recombinant human (rh)PDGF-AB + 20% autologous serum (AS) with and without 10µM AZA for 2 and 23 days respectively (Figure 1A). During daily observations, unilocular lipid globules were observed to fragment within adipocytes ∼ day 10 with progressive extrusion of fat into culture medium, coincident with changes in cell morphology (Movie S1). Consistent with these observations, when fixed and stained with Oil Red O, adipocytes that were globular in shape at the start of culture, resembled lipid laden stromal cells at day 12 and lipid free stromal cells at day 25 (Figure 1B).

**Figure 1.**
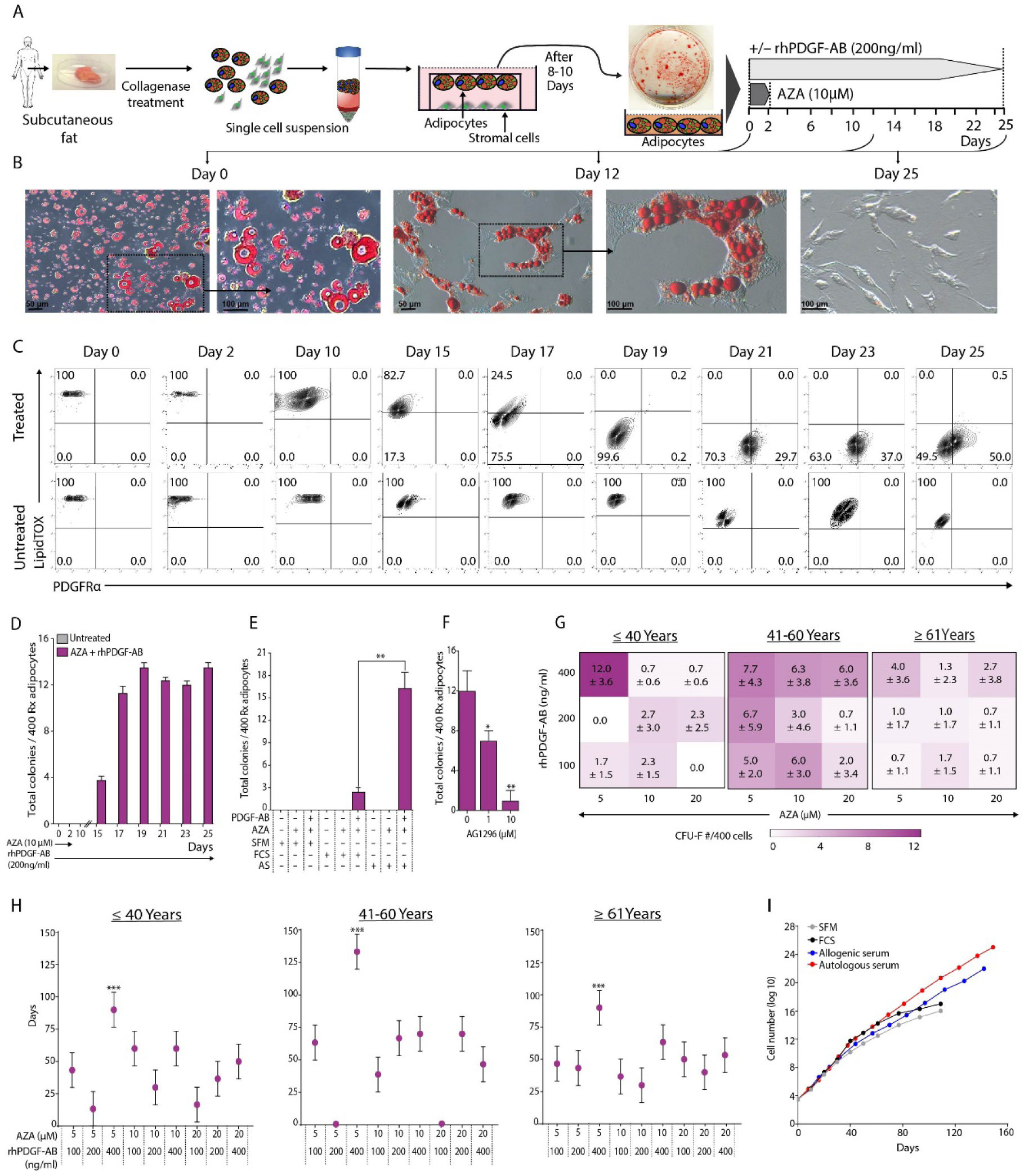
Generating iMS cells from primary human adipocytes. **(A)** Schematic showing steps used to generate plastic-adherent mature adipocytes from donor sub-cutaneous fat, and their subsequent reprogramming. (**B**) Still images (days 0, 12 and 25) of Oil Red O stained adipocytes in medium supplemented with autologous serum following treatment with rhPDGF-AB and AZA as shown in A. (**C**) Flow cytometry plots of LipidTOX (fat) and PDGFRα (mesenchymal) expression in adipocytes cultured as shown in A. (**D**) CFU-F counts from treated and untreated adipocytes harvested at various timepoints during conversion. (**E**) CFU-F counts from treated (Rx) adipocytes (n=400) treated with indicated combinations of rhPDGF-AB, AZA, 20% fetal calf serum (FCS), autologous serum (AS) or serum free media (SFM) as shown in A. (**F**) CFU-F counts from adipocytes reprogrammed as shown in A in the presence of 0, 1μM or 10μM of AG1296, a selective inhibitor of PDGF receptors α and β. (**G**) Heat map representations of CFU-F counts per 400 reprogrammed adipocytes harvested from three donor age groups (age ≤ 40, age 41-60 or age ≥ 61 years; n=3 for each) and generated using indicated dose combinations of rhPDGF-AB and AZA. (**H**) Long term growth potential of reprogrammed adipocytes harvested from three donor age groups (age ≤ 40, age 41-60 or age ≥ 61 years; n=3 for each) and generated using indicated dose combinations of rhPDGF-AB and AZA. (**I**) Long term growth potential of iMS cells cultured in SFM or media supplemented with FCS, autologous or allogeneic serum. rhPDGF-AB; recombinant human platelet derived growth factor AB, AZA; azacitidine, PDGFRα; platelet derived growth factor receptor alpha, CFU-F; colony forming unit-fibroblast, Rx; treated adipocytes, FCS; fetal calf serum, AS; autologous serum, SFM; serum free media, iMS; induced multipotent stem cell. Error bars indicate standard deviation, n =3, *P < 0.05, **P < 0.01, *** P<0.0001 calculated using either a student’s t test (E and F) or a linear mixed model (H).

To evaluate these changes in individual cells, we performed flow cytometry at multiple time points during treatment and probed for adipocyte (LipidTOX) (*24*) and stromal cell characteristics (PDGFRα expression; (*25*)) (Figure 1C). Compared with adipocytes cultured in media supplemented with AS but without PDGF-AB/AZA (Figure 1C lower panel; untreated), a sub-population of cells cultured in the presence of PDGF-AB/AZA (Figure 1C upper panel; treated) showed reduced LipidTOX staining intensity at day 10, with progressive reduction and complete absence in all cells by day 19. Adipocytes expressed PDGFRβ (Figure S1B) but not PDGFRα (Figure 1C) at day 0 but both the frequency and intensity of PDGFRα staining increased from day 21. To record these changes in real-time, we also continuously live-imaged treated adipocytes from day 15 to 25 and recorded the extrusion of fat globules, change in cell morphology from globular to stromal, acquisition of cell motility and cell mitosis (Movie S1 and Figure S1C). CFU-F capacity was absent at day 10, present in day 15 cultures and tripled by day 19 with no substantial increase at days 21, 23 and 25 (Figure 1D). It is noteworthy that CFU-F potential was acquired prior to PDGFRA surface expression when adipocytes had started to display stromal cell morphology and had diminished fat content. There was also no CFU-F capacity in adipocytes cultured in αMEM with fetal calf serum (FCS) or AS, unless supplemented with both PDGF-AB and AZA (Figure 1E). CFU-F capacity was significantly higher with AS than with FCS and absent in serum free media (SFM). As previously shown with reprogramming of murine osteocytes, there was dose dependent inhibition of CFU-F capacity when AG1296, a potent non-selective PDGF receptor tyrosine kinase inhibitor (*26*) was added to the reprogramming media (Figure 1F).

To evaluate the impact of patient age and concentrations of PDGF-AB and AZA on the efficiency of human adipocyte conversion, we harvested sub-cutaneous fat from donors aged ≤ 40 (n=3), 41-60 (n=3) and ≥ 61 (n=3) years and subjected each to three different concentrations of PDGF-AB (100, 200 and 400ng/mL) and three different concentrations of AZA (5, 10 and 20µM) (Figure 1G). Although all combinations supported cell conversion in all donors across the three age groups, 400ng/mL rhPDGF-AB and 5µM AZA yielded the highest number of CFU-Fs (Figure 1G). When these cultures were serially passaged in SFM (with no PDGF-AB/AZA supplementation, which was used for cell conversion only), adipocytes converted with reprogramming media containing 400ng/mL rhPDGF-AB and 5µM AZA, were sustained the longest (Figure 1H and Figure S1D, Table S2). The growth plateau that was observed even with these cultures (i.e. adipocytes converted with 400ng/mL rhPDGF-AB and 5µM AZA when expanded in SFM or FCS), was overcome when cells were expanded in either autologous or allogeneic human serum (Figure 1I). Taken together, these data identify an optimised protocol for converting human primary adipocytes from donors across different age groups and show that these can be maintained long-term in culture.

### *In vitro* characteristics of human iMS cells

The genetic stability of human iMS cells (RM0072 and RM0073) was assessed using SNP arrays and shown to have a normal copy number profile at a resolution of 250kb (Figure S2A). Given the stromal characteristics observed in human adipocytes treated with PDGF-AB/AZA (Figure 1), we performed flow cytometry to evaluate their expression of MSC markers CD73, CD90, CD105 and STRO1 (*13*), and noted expression levels comparable to AdMSCs extracted from the same subcutaneous fat harvest (Figure 2A). The global transcriptomes of iMS cells and matched AdMSCs were distinct from untreated control adipocytes but were broadly related to each other (Figure 2B; (i)-(ii)). Ingenuity pathway analysis (IPA) using genes that were differentially expressed between AdMSCs vs. adipocytes (3307 UP/4351 DOWN in AdMSCs vs. adipoctyes; FDR ≤ 0.05) and iMS vs adipocytes (3311 UP/4400 DOWN in iMS vs. adipoctyes; FDR ≤ 0.05) showed changes associated with gene expression, post-translational modification and cell survival pathways and organismal survival and systems development (Figure 2B (iii)). The number of differentially expressed genes between iMS cells and AdMSCs were limited (2 UP/26 DOWN in iMS vs. AdMSCs; FDR ≤ 0.05) and too few for confident IPA annotation. All differentially expressed genes and IPA annotations are shown in Tables S3 and S4 respectively.

**Figure 2.**
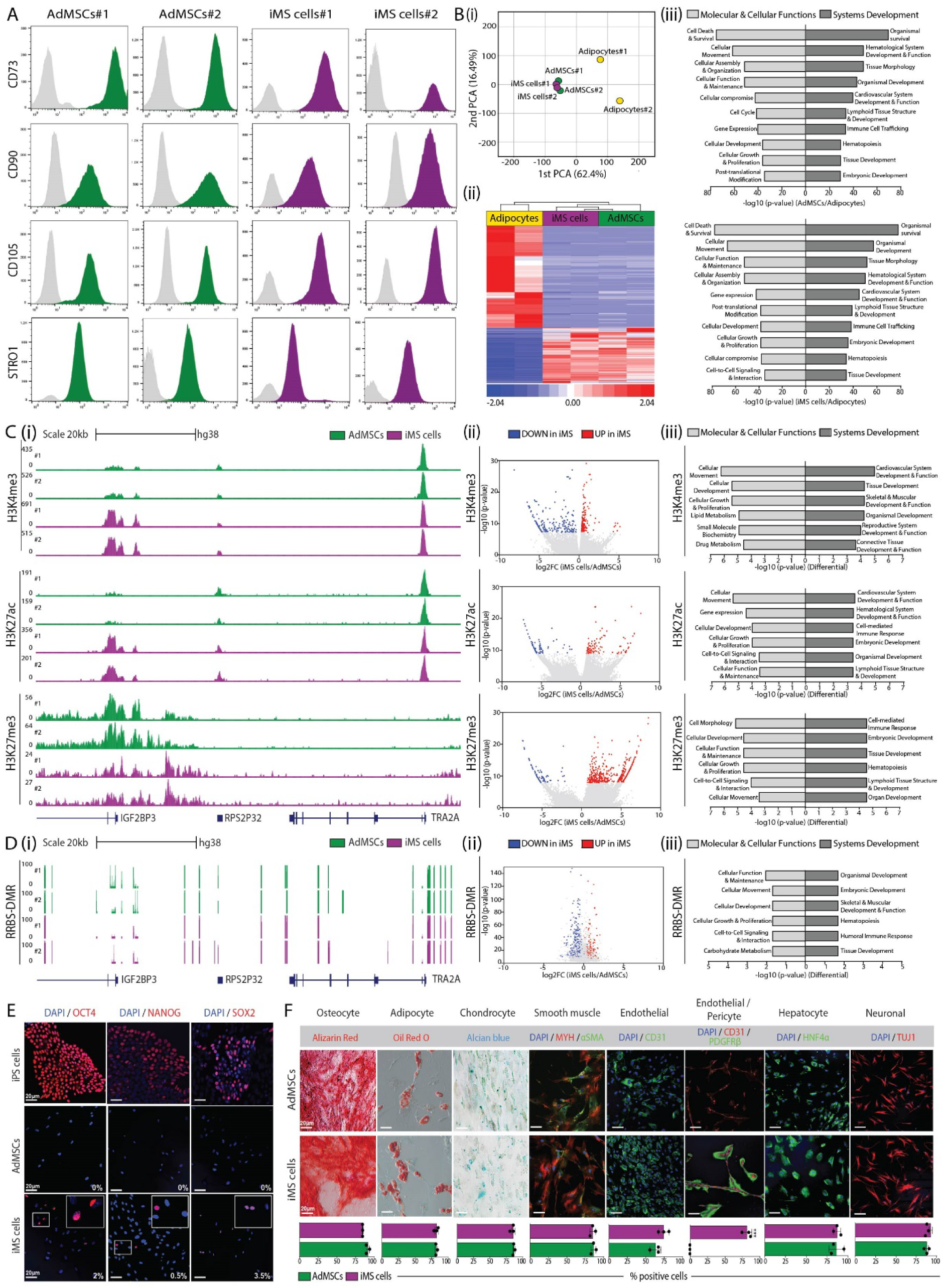
*In vitro* characterisation of human iMS cells. **(A)** Flow cytometry histograms showing expression intensities of mesenchymal stromal cell markers on adipose tissue derived mesenchymal stem cells (AdMSCs; green) and iMS cells (purple) from matched donors (#1; RM0058 and #2; RM0059, passage (P) 2). (**B)** (i) Principal component analysis (PCA) of transcriptomes (RNA-seq) of matched adipocytes, AdMSCs and iMS cells (#1; RM0058 and #2; RM0059). (ii) Hierarchical clustering of genes that were differentially expressed (FDR ≤ 0.05) between primary human adipocytes, AdMSCs and iMS cells. (iii) Ingenuity pathway analysis (IPA) of genes that were differentially expressed between AdMSCs and adipocytes (top) or iMS cells and adipocytes (bottom). The most highly enriched annotated biological functions corresponding to these gene lists are shown. (**C**) (i) H3K4me3, H3K27ac and H3K27me3 ChIP-seq profiles in AdMSCs (green) and iMS cells (purple) from matched donors (#1; RM0057 and #2; RM0072) at a representative locus with differential enrichment. The region shown is chr7:23454137-23538991 (hg38). (ii) Volcano plots of H3K4me3 (top), H3K27Ac (middle) and H3K27me3 (bottom) enrichment peaks that were significantly UP (red) or DOWN (blue) in iMS cells vs. AdMSCs. (iii) IPA using genes corresponding to H3K4me3 (top), H3K27Ac (middle) and H3K27me3 (bottom) peaks that were significantly UP or DOWN in iMS cells vs. AdMSCs. The most highly enriched biological functions corresponding to each gene list is shown. **(D)** (i) DNA methylation levels at a representative region in AdMSCs (green) and iMS cells (purple) from matched donors (#1; RM0057 and #2; RM0072). The region shown is chr7:23454137-23538991 (hg38). (ii) A volcano plot of regions with significantly higher (red) or lower (blue) DNA methylation in iMS cells vs. AdMSCs. (iii) IPA using genes corresponding to differentially methylated regions (DMRs). The most highly enriched biological functions corresponding to this gene list is shown. (**E**) Confocal microscopy images showing OCT4 (left), NANOG (middle), and SOX2 (right) expression in iPS cells (top row), AdMSCs (middle row) and iMS cells (bottom row). The fraction of AdMSCs and iMS cells expressing each protein is listed. **(F)** Microscopy images of AdMSCs (upper panel) and iMS cells (lower panel) differentiated *in vitro* into various lineages and stained for representative proteins. Bar graphs showing staining frequencies (% of cells positive) are shown below each representative image. Error bars indicate standard deviation, n=3. ***P<0.001 (student’s t test). MYH; myosin heavy chain, αSMA; alpha smooth muscle actin, PDGFRβ; platelet derived growth factor receptor beta, HNF4α; Hepatocyte Nuclear Factor 4 Alpha, TUJ1; neuron-specific class III beta-tubulin.

In the absence of significant basal differences in the transcriptomes of AdMSCs and iMS cells, and the use of a hypomethylating agent to induce adipocyte conversion into iMS cells, we examined global enrichment profiles of histone marks associated with transcriptionally active (H3K4me3, H3K27Ac) and inactive (H3K27me3) chromatin. There were differences in enrichment of specific histone marks in matched AdMSCs vs. iMS cells at gene promoters and distal regulatory regions (Figure 2C (i), Figure S2B-D). H3K4me3, H3K27ac and H3K27me3 enrichments were significantly higher at 255, 107 and 549 regions and significantly lower at 222, 78 and 98 regions in iMS cells vs. AdMSCs (Figure 2C (ii); Table S5) and were assigned to 237, 84 and 350 and 191, 58 and 67 genes respectively. IPA was performed using these gene lists to identify biological functions that may be primed in iMS cells relative to AdMSCs (Figure 2C (iii); Table S6). Amongst these biological functions, annotations for molecular and cellular function (cellular movement, development, growth and proliferation) and systems development (general; embryonic and tissue development and specific; cardiovascular, skeletal and muscular and hematological) featured strongly and overlapped across the different epigenetic marks.

We extended these analyses to also assess global CpG methylation in matched AdMSCs and iMS cells using reduced representation bisulfite sequencing (RRBS; (*27*)). Again, there were loci with differentially methylated regions in iMS cells vs. AdMSCs (Figure 2D (i)) with increased methylation at 158 and reduced methylation at 397 regions amongst all regions assessed (Figure 2D (ii)). IPA of genes associated with these DMRs, showed a notable overlap in annotated biological functions (Figure 2D (iii)) with those associated with differential H3K4me3, H3K27Ac and H3K27me3 enrichment (Figure 2C (iii)). Taken together, these data imply that although basal transcriptomic differences between iMS cells and AdMSCs were limited, there were notable differences in epigenetic profiles at *cis*-regulatory regions of genes that were associated with cellular growth and systems development.

PDGF-AB/AZA treated murine osteocytes (murine iMS cells), but not bone derived MSCs, expressed pluripotency associated genes, which were detectable by immunohistochemistry in 1-4% of cells (*14*). To evaluate expression in reprogrammed human cells, PDGF-AB/AZA treated human adipocytes and matched AdMSCs were stained for OCT4, NANOG and SOX2 with expression noted in 2%, 0.5% and 3.5% of iMS cells respectively, but no expression detected in AdMSCs (Figure 2E). MSCs can be induced to differentiate *in vitro* into various cell lineages in response to specific cytokines and culture conditions. To evaluate the *in vitro* plasticity of human iMS cells, we induced their differentiation along with matched AdMSCs, into bone, fat and cartilage, as well as into other mesodermal (Matrigel tube forming assays for endothelial cells (CD31) and pericytes (PDGFRβ) and muscle (MHC, αSMA)), neuroectodermal (neuronal; TUJ1) and endodermal (hepatocyte; HNF4α) lineages (Figure 2F). The differentiation potential of iMS cells and AdMSCs was comparable with the notable exception that only iMS cells generated pericyte lined endothelial tubes in Matrigel. Taken together with the notable differences in epigenetic profiles, these functional differences and low-level expression of pluripotency genes in iMS cell subsets, suggested that these cells could be more amenable than matched AdMSCs to respond to developmental cues *in vivo*.

### *In vivo* characteristics of human iMS cells

To evaluate spontaneous teratoma formation and *in vivo* plasticity of iMS cells, we tagged these cells and their matched AdMSCs with a dual lentiviral reporter, LeGO-iG2-Luc2 (*28*) that expresses both green fluorescent protein (GFP) and luciferase under the control of the cytomegalovirus promoter (Figure 3A). To test teratoma initiating capacity, we implanted tagged cells under the right kidney capsules of NSG mice (n=3/treatment group) after confirming luciferase/GFP expression in cells in culture (Figure S3A and S3B). Weekly bioluminescence imaging (BLI) confirmed retention of cells *in situ* (Figure 3B (i)) with progressive reduction of signal over time (Figure 3B (ii)) and the absence of teratomas in kidneys injected with either AdMSCs or iMS cells (Figure 3B (iii)).

**Figure 3.**
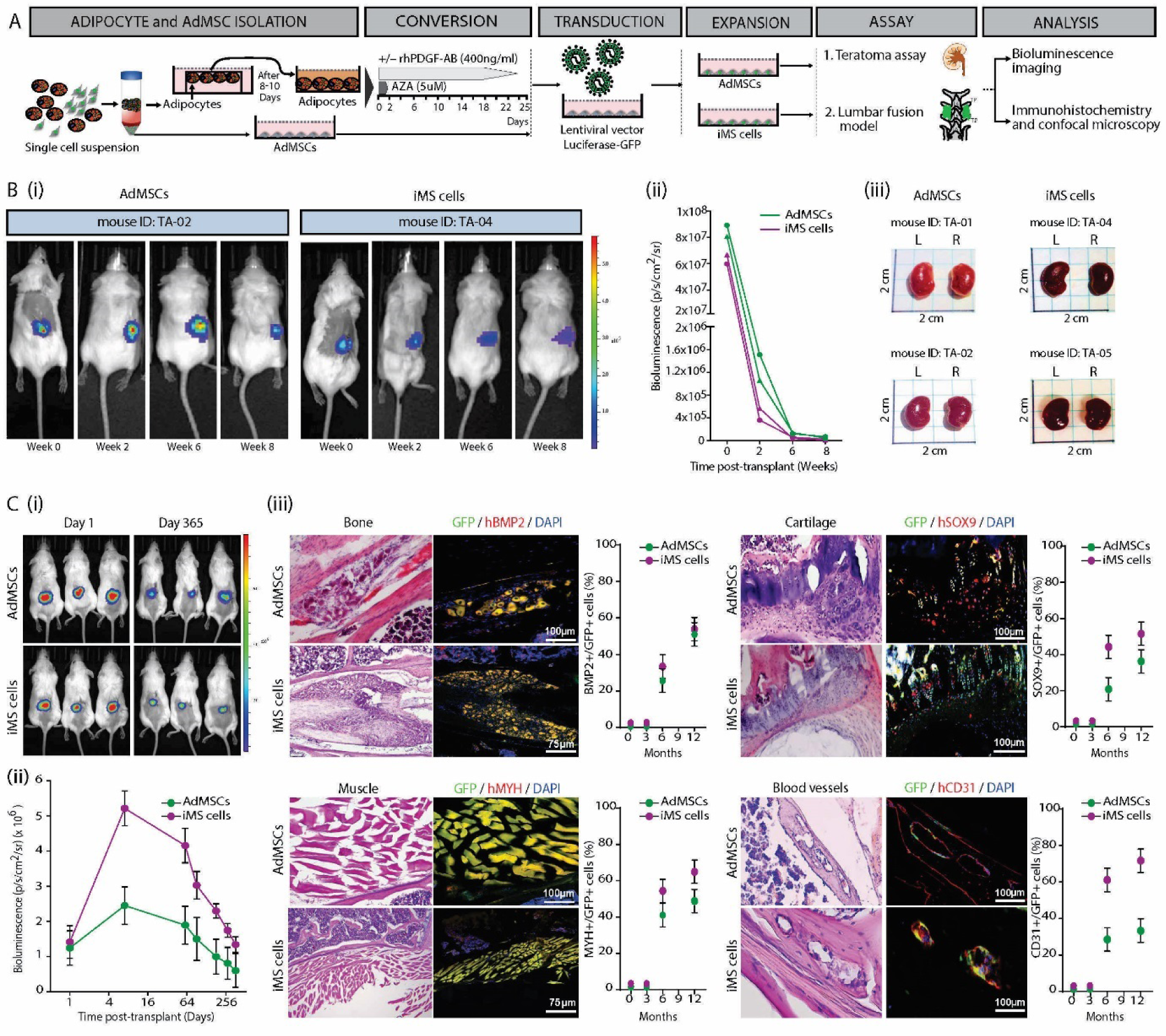
*In vivo* characterisation of human iMS cells. **(A)** Schematic showing generation of Luciferase/GFP-reporter AdMSCs and iMS cells, and downstream assessment of their *in vivo* function. (**B**) Assessment of teratoma initiating capacity; (i) Bioluminescence images of mice taken at 0, 2, 6 and 8 weeks following the implantation of 1 × 10^6 cells matched AdMSCs and iMS cells (P2; RM0057; n=2/group) under the right kidney capsules. (ii) Quantification of bioluminescence over time. (iii) Gross morphology of kidneys following sub-capsular implantation of cells (right) or vehicle control (left) and harvested at 8-weeks post-implantation. (**C**) Assessment of *in vivo* plasticity in a posterior–lateral inter transverse lumbar fusion model; (i) Bioluminescence images of mice following lumbar implantation of 1 × 10^6 matched AdMSCs or iMS cells (P2; RM#38; n=3/group) at 1 day and 365 days post-transplant. (ii) Quantification of bioluminescence over time. (iii) Each panel in the series (bone, cartilage, muscle and blood vessels) shows light microscopy images of tissues harvested at 6-months post either AdMSCs or iMS cells implantation and stained with hematoxylin and eosin (left), and confocal microscopy images of an adjacent tissue section stained with lineage specific anti-human fluorescent antibodies (right). Corresponding graphs show donor cell (GFP^+^) contributions to bone, cartilage, muscle, and blood vessels as a fraction of total (DAPI^+^) cells in 4-5 serial tissue sections. Bars indicate confidence interval, n=3.

To evaluate whether iMS cells survived and integrated with damaged tissues *in vivo*, we implanted transduced human iMS cells and matched AdMSCs controls into a posterior– lateral intertransverse lumbar fusion mouse model (*29*) (Figure 3A). Cells were loaded into Helistat collagen sponges 24 hours prior to implantation into the posterior-lateral gutters adjacent to decorticated lumbar vertebrae of NSG mice (iMS n=9 and AdMSC n=9). Cell retention *in situ* was confirmed by intraperitoneal injection of D-luciferin (150mg/mL) followed by BLI 24 hours post cell implantation, then weekly for the first 6-weeks and monthly up to 12 months from implantation (Figure 3C (i)). The BLI signal gradually decreased with time but persisted at the site of implantation at 12 months, the final assessment time point (Figure 3C (ii)). Groups of mice (n=3 iMS and n=3 AdMSC) were sacrificed at 3, 6 and 12 months and tissues harvested from sites of cell implantation for histology and immunohistochemistry (Figure 3C (iii)). Although implanted iMS cells and AdMSCs were present and viable at sites of implantation at 3-months, there was no evidence of lineage specific gene expression in donor human cells (Figure S3C). By contrast, at 6-months post-implantation, GFP^+^ donor iMS cells and AdMSCs were shown to contribute to new bone (BMP2), cartilage (SOX9), muscle (MYH) and endothelium (CD31) at these sites of tissue injury (Figure 3C (iii)). The proportion of donor cells expressing lineage specific markers in a corresponding tissue section was significantly higher in iMS cells compared with matched AdMSCs at 6 months (Figure 3C (iii), Table S7) as well as 12 months (Figure S3D, Table S7). There was no evidence of malignant growth in any of the tissue sections or evidence of circulating implanted GFP^+^ iMS cells or AdMSCs (Figure S3E).

Taken together, these data show that implanted iMS cells were not teratogenic, were retained long-term at sites of implantation, and contributed to regenerating tissues in a context dependent manner with greater efficiency than matched AdMSCs.

### Regenerative potential of human iMS cells in a skeletal muscle injury model

Although appropriate to assess *in vivo* plasticity and teratogenicity of implanted cells, the posterior–lateral intertransverse lumber fusion mouse model is not suited to address the question of tissue specific differentiation and repair *in vivo*. To this end, we used a muscle injury model (*30*) where necrosis was induced by injecting 10µM cardiotoxin (CTX) into the left tibialis anterior (TA) muscle of 3-month old female SCID/Beige mice. CTX is a myonecrotic agent that spares muscle satellite cells and is amenable to the study of skeletal muscle regeneration. At 24-hours post injury, Matrigel mixed with either 1 × 10^6 iMS cells or matched AdMSCs (or no cells as a control), was injected into the damaged TA muscle. The left (injured) and right (uninjured control) TA muscles were harvested at 1, 2 or 4 weeks post-injury to assess the ability of donor cells to survive and contribute to muscle regeneration without ectopic tissue formation (Figure 4A; Cohort A). Donor human iMS cells or AdMSCs compete with resident murine muscle satellite cells to regenerate muscle, and their regenerative capacity is expected to be handicapped not only by the species barrier but also by having to undergo muscle satellite cell commitment prior to productive myogenesis. Recognising this, a cohort of mice were subject to a second CTX injection, 4 weeks from the first injury/cell implantation followed by TA muscle harvest 4 weeks later (Figure 4A; Cohort B).

**Figure 4.**
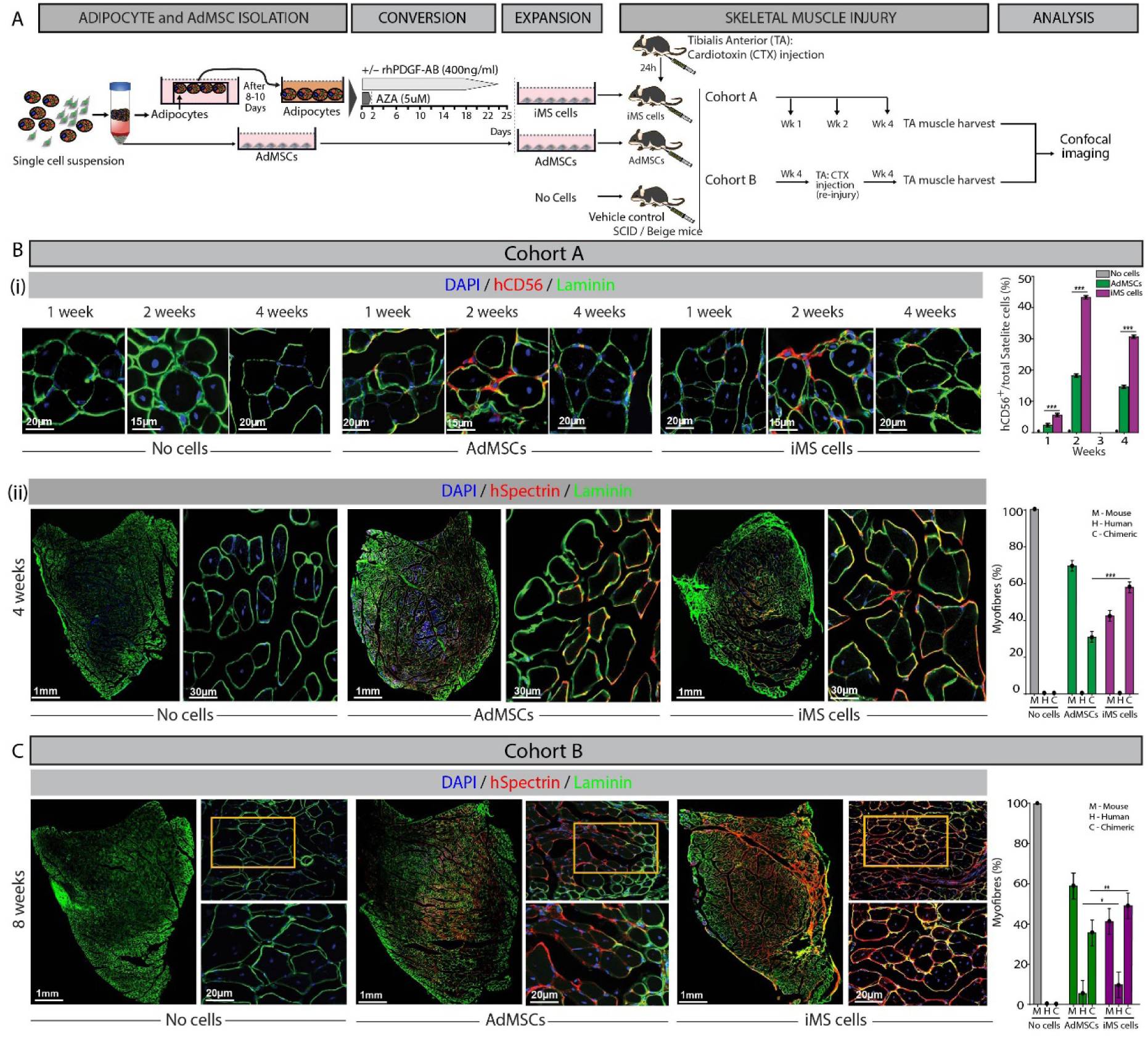
Regenerative potential of human iMS cells in a skeletal muscle injury model. **(A)** Schematic showing steps followed to generate iMS and AdMSCs and to assess their differentiation potential in a cardiotoxin mediated tibialis anterior (TA) muscle injury model. In cohort A, cardiotoxin (CTX; 10µM) was delivered by intramuscular (IM) injection into the left tibialis anterior muscle 24 hours prior to IM injection of 1 × 10^6 matched AdMSCs or iMS cells (P2; RM0057) or Matrigel (vehicle control). TA muscles were harvested at 1, 2 or 4 weeks (n=3 per treatment group at each time point) for histology and confocal microscopy. In cohort B, a second CTX (10µM) injection was delivered to the previously injured and treated muscle 4 weeks following the first injection. TA muscles were harvested 4-weeks later for histology and confocal microscopy (n=3 per treatment group). **(B)** (i) Representative confocal images of TA muscle sections harvested from cohort A at 1, 2, and 4 weeks following injury/treatment and stained for CD56^+^ satellite cells (red; human specific) and Laminin basement membrane protein (green; mouse/human). The graph to the right shows donor hCD56^+^ satellite cell fraction for each treatment group over time. (ii) Confocal images of TA muscle sections harvested at 4 weeks and stained for Spectrin (red; human specific) and Laminin (green; mouse/human). For each treatment, the left panel shows a tile scan of the TA muscle, and the right panel shows a high magnification confocal image. The graph to the right shows the contribution of mouse (M), human (H) or chimeric (C) myofibres (based on hSpectrin^+^/Laminin^+^) in 3-5 serial TA muscle sections/mouse and n= 3 mice per treatment group). **(C)** Representative confocal images of TA muscle sections from cohort B, harvested 4-weeks following re-injury with CTX and stained for Spectrin (red; human specific) and Laminin (green; mouse/human). For each treatment, the left panel shows a tile scan of the TA muscle, the upper right panel shows a low magnification image, and the lower right panel shows a high magnification image of the area boxed above. The graph to the right shows the contribution of mouse (M), human (H) or chimeric (C) myofibres (based on hSpectrin^+^/Laminin^+^) in 3-5 serial TA muscle sections/mouse and n= 3 mice per treatment group). For all graphs, bars indicate confidence interval. *P<0.05, **P<0.01, ***P<0.001 (linear mixed model).

In tissue sections harvested from Cohort A, donor-derived muscle satellite cells (*31*) (hCD56^+^; red) were evident in muscles implanted with both iMS cells and AdMSCs at each time point but were most numerous at 2-weeks post implantation (Figure 4B (i)). The frequency of hCD56^+^ cells relative to total satellite cells (sub-laminar DAPI^+^ cells) was quantified in 3-5 serial sections of TA muscles per mouse in each of three mice per treatment group, and was noted to be higher following the implantation of iMS cells compared with AdMSCs at all time points (week 1; 5.6% vs 2.4%, week 2; 43.3% vs. 18.2% and week 4; 30.7% vs. 14.6%; Figure 4B (i); Table S8). Donor cell contribution to regenerating muscle fibres was also assessed by measuring human spectrin (*32*) co-staining with mouse/human laminin (*33*) at 4 weeks (Figure 4B (ii)). At least 1000 myofibres from 3-5 serial sections of TA muscles for each of three mice in each treatment group were scored for human (H; hSpectrin^+^ (full circumference); laminin^+^), murine (M; mouse; hSpectrin^-^; laminin^+^) or mouse/human chimeric (C; hSpectrin+ (partial circumference); laminin^+^) myofibers. Although none of the myofibres seen in cross section appeared to be completely human (i.e. donor-derived), both iMS cells and AdMSCs contributed to chimeric myofibres (Figure 4B (ii)). iMS cell implants contributed to a substantially higher proportion of chimeric fibres than AdMSC implants (57.7% vs 30.7%: Table S9). In Cohort B, TA muscles were allowed to regenerate following the initial CTX injection/cell implantation, and re-injured 4-weeks later with a repeat CTX injection. In these mice, although total donor cell contributions to myofibres in TA muscles harvested 4-weeks post re-injury were comparable to that observed in Cohort A, there were now myofibres that appeared to be completely human (Figure 4C). There were substantially more human myofibres following iMS cell implants than with AdMSCs (9.7% vs. 5.4%; Table S10). There was no evidence of ectopic tissue formation in TA muscles following implantation of either iMS cells or AdMSCs in either cohort. Taken together, these data showed that iMS cells had the capacity to respond appropriately to the injured environment and contribute to tissue specific regeneration.

## DISCUSSION

We have optimised a protocol, originally designed for mouse osteocytes, to convert human primary adipocytes into iMS cells. We show that these long-term repopulating cells regenerate tissues *in vivo* in a context dependent manner without generating ectopic tissues or teratomas.

PDGF-AB, AZA and serum are indispensable ingredients in reprograming media but how they cooperate to induce cell conversion is speculative. PDGF-AB is reported to bind and signal via PDGFR-αα and -αβ but not -ββ subunits (*21*). Mouse osteocytes and human adipocytes lack PDGFRα, although surface expression was detectable as cells transition during reprogramming (mouse; day 2/8 (*14*) and human day 21/25). However, both mouse osteocytes and human adipocytes express PDGFRβ (*14*). Given that PDGFR inhibition attenuates iMS cell production in both mice (*14*) and humans a degree of facilitated binding of PDGF-AB to - ββ subunits or signaling through a non-canonical receptor is likely to occur, at least at the start of reprogramming.

PDGF-AB was replenished in culture throughout the reprogramming period but AZA treatment was limited to the first two days for both mouse osteocyte and human adipocyte cultures. DNA replication is required for incorporation of AZA into DNA (*34*) and as such DNA demethylation is unlikely to be an initiating event in the conversion of terminally differentiated non-proliferating cells such as osteocytes and mature adipocytes. However, the majority of intracellular AZA is incorporated into RNA, which could directly impact the cellular transcriptome and proteome as an early event (*35, 36*). It is feasible that subsequent redistribution of AZA from RNA to DNA occurs when cells replicate resulting in DNA hypomethylation as a later event (*37*).

In the absence of serum, we could neither convert primary human adipocytes into iMS cells nor perpetuate these cells long-term in culture. The efficiency of conversion and expansion was significantly higher with human vs. fetal calf serum and highest with autologous serum. The precise serum factor(s) that are required for cell conversion in conjunction with PDGF-AB and AZA are not known. The volumes of blood (∼ 50mLs x 2) and sub-cutaneous fat (5g) that we harvested from donors were not limiting to generate sufficient numbers of P2 iMS cells (∼10 × 10^6) for *in vivo* implantation and are in the range of cell numbers used in prospective clinical trials using mesenchymal precursor cells for chronic discogenic lumbar back pain (NCT02412735; 6 × 10^6) and hypoplastic left heart syndrome (NCT03079401; 20 × 10^6).

Producing clinical-grade autologous cells for cell therapy is expensive and challenging requiring suitable quality control measures and certification. However, the advent of chimeric antigen receptor-T cell therapy into clinical practice (*38*), has shown that production of a commercially viable, engineered autologous cellular product is feasible where a need exists. Although there were no apparent genotoxic events in iMS cells at P2, *ex vivo* expansion of cells could risk accumulation of such events and long-term follow up of on-going and recently concluded clinical trials using allogeneic expanded mesenchymal progenitor cells will be instructive with regards to their teratogenic potential. The expression of pluripotency factors in 2-3% of murine and human iMS cells did not confer teratogenic potential in teratoma assays or at 12-months follow-up despite persistence of cells at the site of implantation. However, this risk cannot be completely discounted and the clinical indications for iMS or any cell therapy requires careful evaluation of need.

In regenerating muscle fibres, it was noteworthy that iMS cells appeared to follow canonical developmental pathways in generating muscle satellite cells that were retained and primed to regenerate muscle following a second muscle specific injury. Although iMS cells were generated from adipocytes, there was no evidence of any adipose tissue generation. This supports the notion that these cells have lost their native differentiation trajectory and adopted an epigenetic state that favoured response to local differentiation cues. The superior *in vivo* differentiation potential of iMS cells vis a vis matched AdMSCs was consistent with our data showing that the epigenetic state of iMS cells was better primed for systems development. Another clear distinction between iMS cells and AdMSCs was the ability of the former to produce CD31^+^ endothelial tube-like structures that were enveloped by PDGFRβ^+^ pericytes. An obvious therapeutic application for iMS cells in this context is vascular regeneration in the setting of critical limb ischemia to restore tissue perfusion, an area of clear unmet need (*39*).

An alternative to *ex vivo* iMS cell production and expansion is the prospect of *in situ* reprogramming by local sub-cutaneous administration of the relevant factors to directly convert sub-cutaneous adipocytes into iMS cells, thereby eliminating the need for *ex vivo* cell production. AZA is used in clinical practice and administered as a daily sub-cutaneous injection for up to seven days in a 28-day cycle with responders occasionally remaining on treatment for decades (*40*). Having determined the optimal dose of AZA required to convert human adipocytes into iMS cells *in vitro* (2 days, 5µM), the bridge to ascertaining the comparable *in vivo* dose would be to first measure levels of AZA incorporation in RNA/DNA following *in vitro* administration and match the dose of AZA to achieve comparable tissue levels *in vivo*. AZA-MS, a mass spectrometry-based assay was developed to measure *in vivo* incorporation of AZA metabolites in RNA/DNA and is ideally suited to this application (*37*). The duration of AZA administration for adipocyte conversion was relatively short (i.e. 2 days), but PDGF-AB levels were maintained for 25 days. One mechanism of potentially maintaining local tissue concentrations would be to engineer growth factors to bind extra cellular matrices and be retained at the site of injection. VEGF-A and PDGF-BB have recently been engineered with enhanced syndecan binding and shown to promote tissue healing (*41*). A comparable approach could help retain PDGF-AB at the site of injection and maintain local concentrations at the required dose. Whilst our current data show that human adipocyte derived iMS cells regenerate tissues in a context dependent manner without ectopic or neoplastic growth, these approaches are worth considering as an alternative to an *ex vivo* expanded cell source in the future.

## MATERIALS AND METHODS

Extended methods for cell growth and differentiation assays and animal models are available in Supplementary Figures, Tables and Methods, and antibodies used are detailed in Table S11.

### Study design

The primary objective of this study was to optimise conditions that were free of animal products for the generation of human iMS cells from primary adipocytes and to characterise their molecular landscape and function. To this end, we harvested sub-cutaneous fat from donors across a broad age spectrum and used multiple dose combinations of a recombinant human growth factors and a hypomethylating agent used in the clinic and various serum types. We were particularly keen to demonstrate cell conversion and did so by live imaging and periodic flow cytometry for single cell quantification of lipid loss and gain of stromal markers. Using our previous report generating mouse iMS cells from osteocytes and adipocytes as a reference, we first characterised the *in vitro* properties of human iMS cells including (i) long-term growth (ii) colony forming potential (iii) in vitro differentiation and (iv) molecular landscape. Consistent with their comparative morphology, cell surface markers and behavioural properties, the transcriptomes (RNA-seq) were broadly comparable between iMS cells and matched AdMSCs leading to investigation of epigenetic differences (ATAC-seq, Histone ChIP-seq and RRBS for DNA methylation differences) that might explain properties that were unique to iMS cells (expression of pluripotency factors, generation of endothelial tubes *in vitro* with pericyte envelopes and in vivo regenerative potential). Context dependent *in vivo* plasticity was assessed using a tissue injury model that was designed to promote bone/cartilage/muscle/blood vessel contributions from donor cells and simultaneously assess the absence of ectopic/malignant tissue formation by these cells (labelled and tracked *in vivo* using a bioluminescence/fluorescence marker). Tissue specific regeneration and the deployment of canonical developmental pathways was assessed using a specific muscle injury model and donor cell contributions in all injury models was performed on multiple serial tissue sections in multiple mice with robust statistical analyses (see below). Power calculations were not used, samples were not excluded, and investigators were not blinded. Experiments were repeated multiple times or assessments were performed at multiple time points. Cytogenetic and CNV analyses were performed on iMS and AdMSCs pre-transplant and their teratogenic potential was assessed both by specific teratoma assays and long-term implantation studies.

### Tissue harvest and cell isolation

Sub-cutaneous fat and blood was harvested from patients undergoing surgery at the Prince of Wales Hospital, Sydney. Patient tissue was collected in accordance with NHMRC National Statement on Ethical Conduct in Human Research (2007) and with approval from the South Eastern Sydney Local Health District Human Research Ethics Committee (HREC 14/119). Adipocytes were harvested as described (*42*). Briefly, adipose tissue was minced and digested with 0.2% Collagenase type 1 (Sigma) at 37°C for 40 minutes and the homogenised suspension passed through a 70µm filter, inactivated with autologous serum and centrifuged. Primary adipocytes from the uppermost fatty layer were cultured using the ceiling culture method (*43*) for 8-10 days. AdMSCs from the stromal vascular pellet were cultured in αMEM+20% autologous serum +100 μg/mL penicillin and 250 ng/mL streptomycin, 200 mM L-Glutamine (complete medium).

### Cell reprogramming

Mature adipocytes were cultured in complete medium supplemented with AZA (R&D Systems; 5, 10, 20µM; 2 days) and rhPDGF-AB (Miltenyi Biotec; 100, 200, 400ng/mL; 25 days) with medium changes every 3-4 days. For inhibitor experiments, AG1296 was added for the duration of the culture. Live imaging was performed using an IncuCyte S3 (10x 0.25 NA objective) or a Nikon Eclipse Ti-E (20x 0.45 NA objective). Images were captured every 30mins for a period of 8 days starting from Day 15. 12-bit images were acquired with a 1280×1024 pixel array and analysed using ImageJ software.

### Animals

Animals were housed and bred with approval from the Animal Care and Ethics Committee, UNSW (17/30B, 18/122B, 18/134B). NSG (NOD.*Cg-Prkdc*^*scid*^*Il2rg*^*tm1Wjl*^/SzJ) and SCID/Beige (C.B-*Igh*-1b/GbmsTac-*Prkdc*^*scid*^*-Lyst*^*bg*^ N, sourced from Charles River) strains were used as indicated. The IVIS Spectrum CT (Perkin Elmer) was used to capture bioluminescence. Briefly, 15 minutes after intraperitoneal injection of D-luciferin (150 mg/kg), images were acquired for 5 minutes and radiance (p/s/cm^2^/sr) was used for subsequent data analysis. The scanned images were analysed using the Living Image 5.0 software (Perkin Elmer).

### Teratoma Assay

Teratoma assays (*44*) were performed on 3-4-month-old female NSG mice. 5 × 10^5^ lentiviral tagged cells in 20 µL of PBS containing 80% Matrigel were injected under the right kidney capsule using a fine needle (26G) and followed weekly by bioluminescence imaging till sacrifice at week 8. Both kidneys were collected, fixed in 4% PFA for 48 hours, embedded in OCT, cryosectioned and imaged for GFP.

### Tissue injury models

#### Posterior-lateral intervertebral disc injury model (29)

1×10^6^ lentiviral tagged (*28*) AdMSCs or iMS cells were loaded onto Helistat collagen sponges and implanted into the postero-lateral gutters in the L4/5 lumbar spine region of anesthetised NSG mice following decortication of the transverse processes. Animals were imaged periodically for bioluminescence to track the presence of transplanted cells. At 3, 6 or 12 months, mice were sacrificed and spines from the thoracic to caudal vertebral region, including the pelvis, were removed whole. The specimens were fixed in 4% PFA for 48 hours, decalcified in 14% (w/v) EDTA, and embedded in OCT. 5 µm sagittal cryosections of fixed tissue were used for histology and immunofluorescence.

#### Muscle injury model (45)

The Left Tibialis anterior (TA) muscle of 3-4-month-old female SCID/Beige mice was injured by injection with 15 µL of 10 µM cardiotoxin (Latoxan). Confocal images of 3-4 serial sections/mouse were captured by Zen core/ AxioVision (Carl Zeiss) and visualised by ImageJ with the colocalization and cell counter plugins (NIH; (*46*)).

### RNA sequencing

Total RNA was extracted using the miRNeasy kit (Qiagen) according to manufacturer’s instructions, and 200ng of total RNA used for Illumina TruSeq library construction. Library construction and sequencing was performed by Novogene (HK) Co., Ltd. Raw paired-end reads were aligned to the reference genome (hg19) using STAR (*47*) and HTSeq (*48*) was used to quantify the transcriptomes using the reference “refFlat” database from the UCSC Table Browser (*49*). The resulting Gene Expression Matrix was normalized and subject to differential gene expression using DeSeq2 (*50*). Normalized gene expression was employed to compute and plot two-dimension Principal Component Analysis (PCA), using the python modules sklearn (v0.19.1) (*51*) and matplotlib (v2.2.2) (*52*) respectively. Differentially expressed genes (log2fold-change ≥ |1|, adj-pval < 0.05) were the input to produce an unsupervised hierarchical clustering heatmap in Partek Genomics Suite (PGS) software (version 7.0) (Partek Inc., St. Louis, MO, USA). Raw data is available using accession GSE150720.

### ChIP sequencing

Chromatin immunoprecipitation (ChIP) was performed as previously described (*53*) using antibodies against H3K27Ac, H3K4Me3, and H3K27Me3. Library construction and sequencing was performed by Novogene (HK) Co., Ltd. Paired end reads were aligned to the hg38 genome build using BWA (*54*) duplicate reads removed using Picard (http://broadinstitute.github.io/picard/), and tracks generated using Deeptools bamCoverage (*55*). Peaks were called using MACS2 (*56*) with the parameter (p = 1e-9). Differentially bound regions between the AdMSC and iMS were calculated using DiffBind (*57*) and regions annotated using ChIPseeker (*58*). Raw data is available using accession GSE151527.

### Reduced Representation Bisulphite Sequencing (RRBS)

Total genomic DNA was extracted using DNA MiniPrep kit (Qiagen) and RRBS library construction and sequencing was performed by Novogene (HK) Co., Ltd. Raw RRBS data in fastq format were quality and adapter trimmed using trim_galore (0.6.4) with --rrbs parameter (http://www.bioinformatics.babraham.ac.uk/projects/trim_galore). The trimmed fastq files were then aligned to a bisulfite converted genome (Ensembl GRCh38) using Bismark (2.3.5) and methylation status at each CpG loci were extracted (*59*). The cytosine coverage files were converted to BigWig format for visualisation. Differentially methylated cytosines (DMCs) and differentially methylated regions (DMRs) were identified using methylKit (1.10) and edmr (0.6.4.1) packages in R (3.6.1) (*60, 61*). DMCs and DMRs were annotated using ChIPseeker (*58*) and pathway enrichment was performed as detailed below. Raw data is available using accession GSE151527.

### Pathway Analysis

Ingenuity Pathway Analysis (IPA; Qiagen) was used to investigate enrichment in Molecular and Cellular Functions, Systems Development and Function and Canonical Pathways.

### Statistical analysis

Statistical analysis was performed in SAS. For the dose optimization experiments (Figure 1), a linear mixed model with participant-level random effects was used to estimate maximum time by dose level and age group. A linear mixed model with participant-level random effects was used to analyse statistical differences in lineage contribution outcomes between treatment groups (Figure 3) and at different timepoints post-transplant, to estimate the percentage of cells by treatment and lineage. For the *in vivo* regeneration experiment (Figure 4), a linear model was used to model the percent of cells over time for each group. Quadratic time terms were added to account for the observed increase from 1 to 2 weeks and decrease from 2 to 4 weeks. In the muscle regeneration experiment a linear model was applied to Cohort A and Cohort B, in order to estimate and compare percent cells by treatment and source.

## ACKNOWLEDGEMENTS

The authors are indebted to the patients who kindly donated tissue to this project. The authors thank Dr Erin Cook (Prince of Wales Private Hospital), Brendan Lee (Mark Wainwright Analytical Centre, UNSW Sydney), and technicians at the UNSW BRC Facility for assistance with sample and data collection and animal care, and Dr Ashwin Unnikrishnan and Dr Chris Jolly for helpful discussions and critical reading of the manuscript. The authors acknowledge the facilities and scientific and technical assistance of the National Imaging Facility, a National Collaborative Research Infrastructure Strategy (NCRIS) capability, at the BRIL (UNSW). The STRO-1 antibody was a kind gift from Professor Stan Gronthos, University of Adelaide, Australia.

## Author contributions

The project was conceived by VC and JEP and the study design and experiments were planned by AY, VC and JEP. Most of the experiments and data analyses were performed by AY, guided and supervised by VC and JEP. SS, RO, DC, FY, MLT, TH, PH, WW and VC performed additional experiments and data analyses with further supervision from RM, CP, CL, JAIT, DC, JWHW, LBH, DB and EH. Statistical analyses were performed by JO. RM, DDM, JCDM, JM and MK provided critical reagents. The manuscript was written by AY, JAIT, VC and JEP, and reviewed and agreed to by all co-authors.

