## Supplementary materials for "Induction of Muscle Regenerative Multipotent Stem Cells from Human Adipocytes by PDGF-AB and 5-Azacytidine"

Yeola et al

### Supplementary Figures

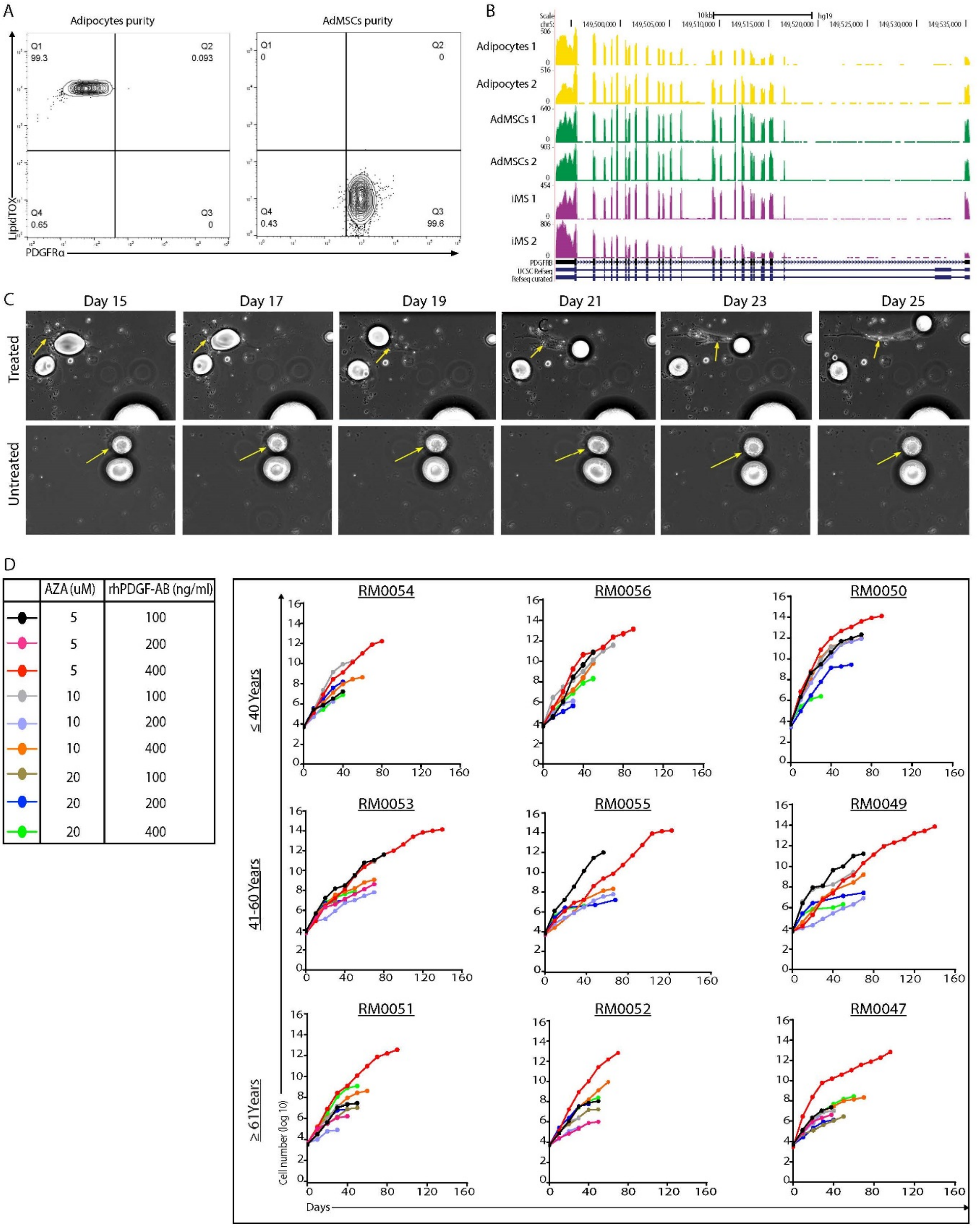


| | AZA (uM) | rhPDGF-AB (ng/ml) |
|---|---|---|
| black | 5 | 100 |
| pink | 5 | 200 |
| red | 5 | 400 |
| grey | 10 | 100 |
| light purple | 10 | 200 |
| orange | 10 | 400 |
| olive | 20 | 100 |
| blue | 20 | 200 |
| green | 20 | 400 |

Figure S1

**Figure S1. (A)** Flow cytometry purity check of freshly isolated primary adipocytes and AdMSCs on the basis of LipidTOX and PDGFRα expression. **(B)** UCSC browser tracks showing *PDGFRB* gene expression in primary adipocytes, AdMSCs and iMS cells converted from primary adipocytes. Region shown is chr5: 149,493,402-149,535,422 (hg19). **(C)** Still images (days 15, 17, 19, 21, 23 and 25) of adipocytes cultured in medium supplemented with autologous serum (AS) following treatment with (upper panel) or without (lower panel) rhPDGF-AB (200ng/mL; 25 days) and AZA (10µM; first 48 hours). Arrows in upper panel indicate changes in cell morphology during reprogramming of an adipocyte into an iMS cell. Arrows in lower panel indicate an adipocyte with unchanged morphology in the absence of rhPDGF-AB and AZA. **(D)** Long-term growth curves of iMS cells converted from primary adipocytes harvested from patients (age groups ≤ 40 years, 41-60 years or ≥ 61 years; 3 each) treated with rhPDGF-AB (100, 200 or 400 ng/mL) and AZA (5, 10 or 20 µM) and expanded in serum free medium (SFM).

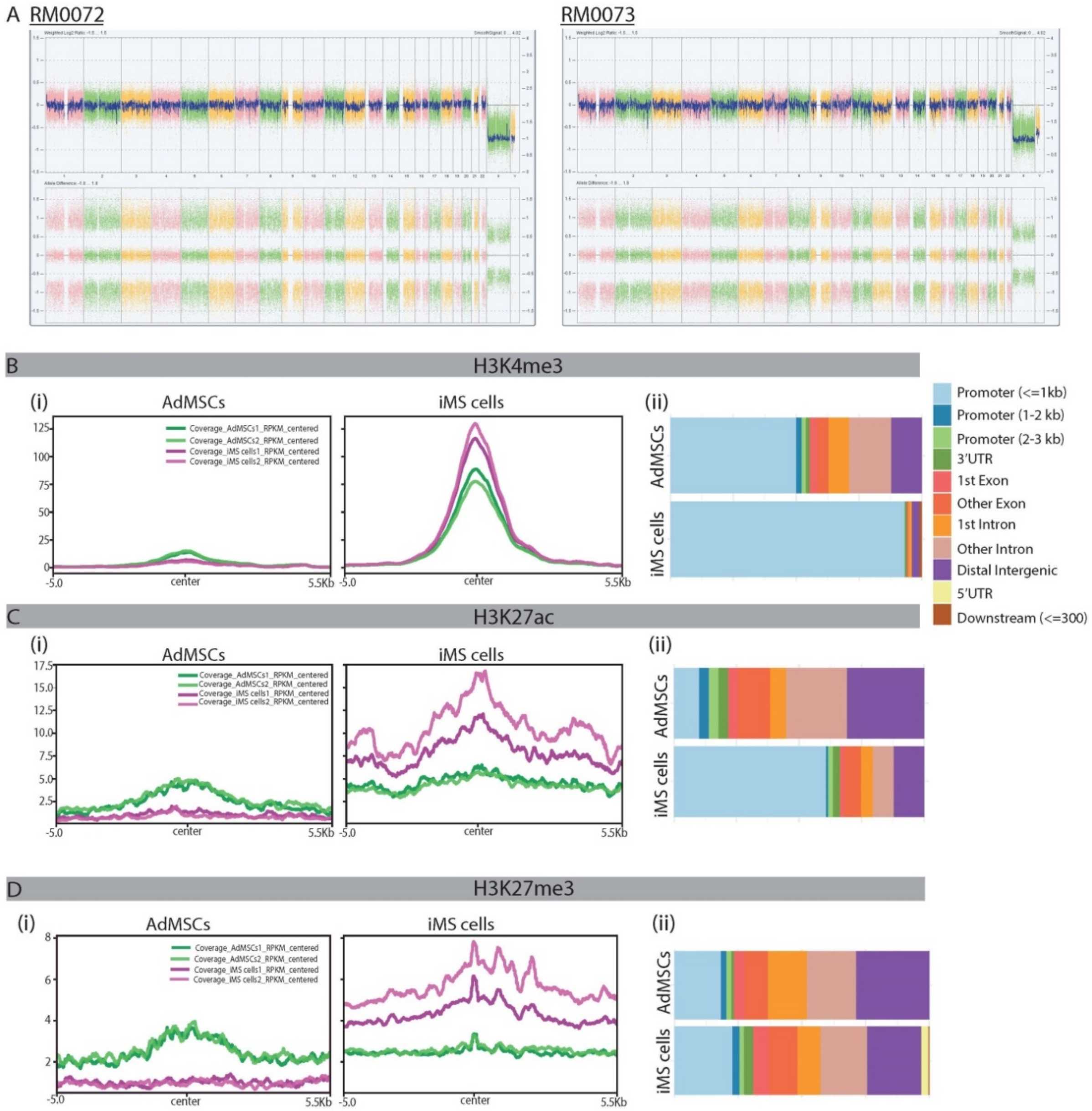
A RM0072
RM0073
B
H3K4me3
(i)
AdMSCs
iMS cells
Coverage_AdMSCs1_RPKM_centered
Coverage_AdMSCs2_RPKM_centered
Coverage_iMS cells1_RPKM_centered
Coverage_iMS cells2_RPKM_centered
125
100
75
50
25
0
-5.0
center
5.5Kb
(ii)
AdMSCs
iMS cells
Promoter (<=1kb)
Promoter (1-2 kb)
Promoter (2-3 kb)
3'UTR
1st Exon
Other Exon
1st Intron
Other Intron
Distal Intergenic
5'UTR
Downstream (<=300)
C
H3K27ac
(i)
AdMSCs
iMS cells
17.5
15.0
12.5
10.0
7.5
5.0
2.5
-5.0
center
5.5Kb
(ii)
AdMSCs
iMS cells
D
H3K27me3
(i)
AdMSCs
iMS cells
8
6
4
2
-5.0
center
5.5Kb
(ii)
AdMSCs
iMS cells

Figure S2

**Figure S2. (A)** SNP analysis for iMS cells cultured for two passages showing that iMS cells derived from two independent patients have normal genome-wide cytogenetic profiles. **(B)** (i) Average histone H3K4me3 profiles around peak centres of regions that are differentially bound in iMS cells as compared to AdMSCs. (ii) Genomic distribution of regions with differential H3K4me3 binding. Plots show regions with increased binding in AdMSC and iMS cells. **(C)** (i) Average histone H3K27ac profiles around peak centres of regions that are differentially bound in iMS cells as compared to AdMSCs. (ii) Genomic distribution of regions with differential H3K27ac binding. Plots show regions with increased binding in AdMSC and iMS cells. **(D)** (i) Average histone H3K27me3 profiles around peak centres of regions that are differentially bound in iMS cells as compared to AdMSCs. (ii) Genomic distribution of regions with differential H3K27me3 binding. Plots show regions with increased binding in AdMSC and iMS cells.

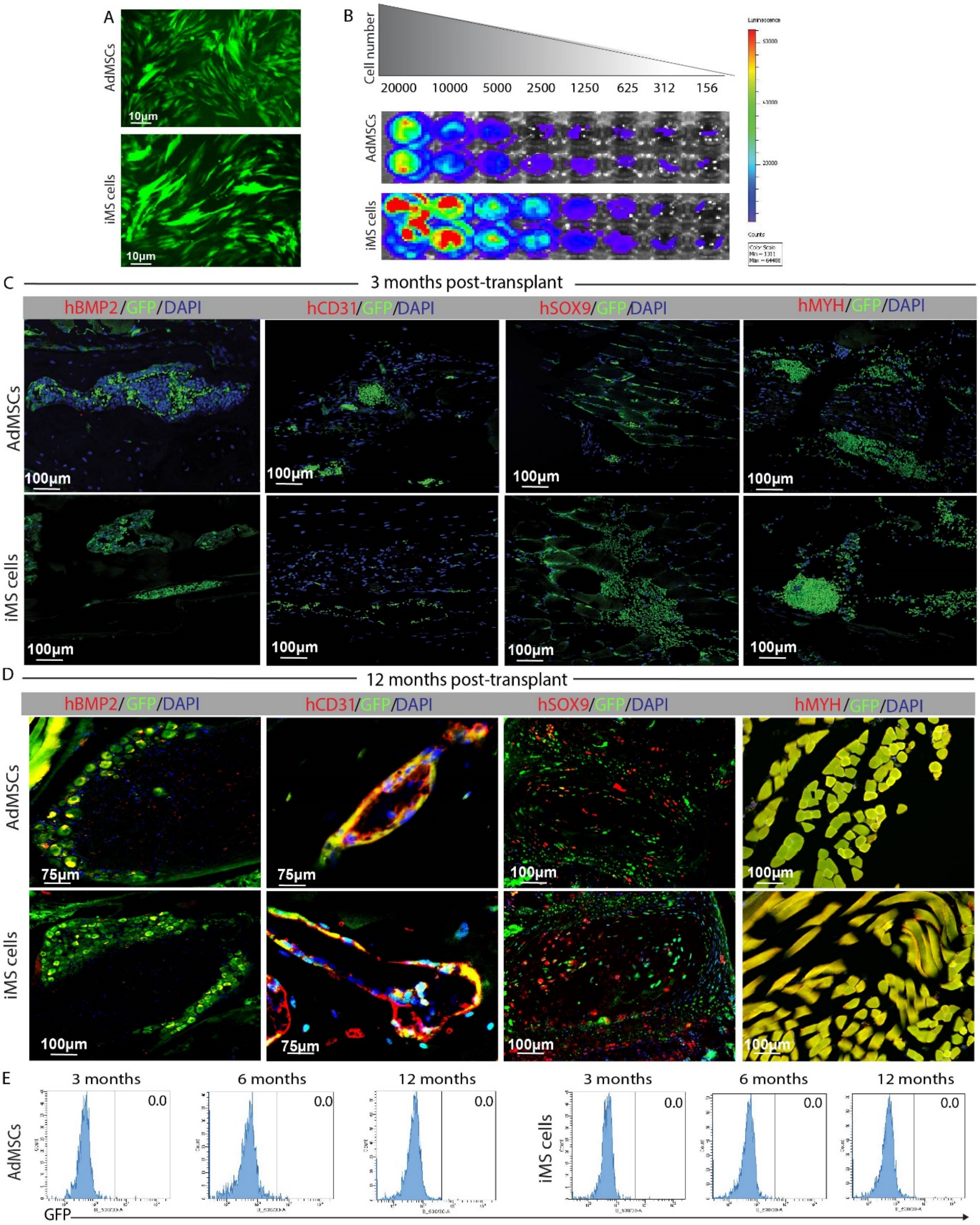
A
AdMSCs
iMS cells
10µm
B
Cell number
20000 10000 5000 2500 1250 625 312 156
C
3 months post-transplant
hBMP2/GFP/DAPI
hCD31/GFP/DAPI
hSOX9/GFP/DAPI
hMYH/GFP/DAPI
100µm
D
12 months post-transplant
75µm
E
3 months
6 months
12 months
0.0
GFP

Figure S3

**Figure S3. (A)** *In vitro* validation of GFP expression in lentiviral tagged cell lines used for *in vivo* safety and efficacy studies. **(B)** *In vitro* validation of bioluminescence in lentiviral tagged cell lines used for *in vivo* safety and efficacy studies. **(C)** Immunofluorescence analysis of tissue sections harvested from spine sites implanted with AdMSCs or iMS cells harvested at 3 months post-transplant (n=3). GFP positive donor cells are maintained at the graft site but remain undifferentiated as assessed by morphology and absence of markers specific for human osteogenic (hBMP2), endothelial (hCD31), chondrogenic (hSOX9), or myogenic (hMYH) lineages. **(D)** Immunofluorescence analysis of tissue sections harvested from spine sites implanted with AdMSCs or iMS cells at 12-months post-transplantation (n=3). Each panel in the series (bone, blood vessels, cartilage, and muscle) shows confocal microscopy images stained with lineage specific anti-human fluorescent antibodies. **(E)** Flow cytometry analysis of GFP positive donor cells in peripheral blood collected from mice at 3, 6, and 12 months post-transplant (n=3 for each time point).

### Supplementary Tables

**Table S1.** Donor details with demographic information for all study participants.

| Patient ID | Age (years) | Gender |
|---|---|---|
| RM0032 | 43 | M |
| RM0035 | 29 | F |
| RM0038 | 40 | F |
| RM0040 | 27 | M |
| RM0042 | 30 | F |
| RM0043 | 33 | M |
| RM0045 | 41 | M |
| RM0047 | 66 | F |
| RM0049 | 47 | F |
| RM0050 | 35 | F |
| RM0051 | 66 | M |
| RM0052 | 65 | M |
| RM0053 | 55 | F |
| RM0054 | 34 | M |
| RM0055 | 45 | M |
| RM0056 | 27 | F |
| RM0057 | 47 | F |
| RM0058 | 49 | M |
| RM0059 | 40 | M |
| RM0061 | 31 | F |
| RM0067 | 32 | M |
| RM0069 | 45 | M |
| RM0071 | 27 | M |
| RM0064 | 43 | M |
| RM0072 | 40 | M |
| RM0073 | 45 | M |

**Table S11**. List of antibodies.

| **Target** | **Dilution** | **Cat. #** | **Supplier** |
|---|---|---|---|
| PDGFRA | 1:100 | 323508 | BioLegend |
| H3K27ac | 5ug/IP | ab4729 | Abcam |
| H3K4Me3 | 5ug/IP | ab8580 | Abcam |
| H3K27Me3 | 5ug/IP | C15410195 | Diagenode |
| STRO1 | 1:50 | N/A | Prof. Stan Gronthos |
| OCT4 | 1:50 | 653701 | Australian Biosearch |
| SOX2 | 1:50 | 656102 | Australian Biosearch |
| Nanog | 1:50 | 674201 | Australian Biosearch |
| Myosin Heavy Chain (MYH) | 1:50 | 621202 | BioLegend |
| α smooth muscle actin (αSMA) | 1:400 | C6198 | Sigma Aldrich |
| CD31 | 1:100 | 303102 | BioLegend |
| PDGFRβ | 1:100 | ab32570 | Abcam |
| HNF4α | 1:100 | ab41898 | Abcam |
| TUJ1 | 1:100 | AB9354 | Millipore |
| BMP2 | 1:100 | sc-6895 | Santacruz Biotechnology |
| SOX9 | 1:150 | sc-17341 | Santacruz Biotechnology |
| Laminin | 1:100 | L9393 | ThermoFisher Scientific |
| CD56 | 1:50 | MA5-11563 | ThermoFisher Scientific |
| Spectrin | 1:60 | NCL-SPEC1 | Novacastra (Leica) |

**Additional Supplementary Tables (excel files)**

Table S2 - Linear mixed model data table used to determine the dose of rhPDGF-AB and AZA for optimal reprogramming of primary adipocytes.

Table S3 - List of differentially expressed genes

Table S4 - List of IPA hits for differentially expressed genes

Table S5 - List of differentially bound regions and differentially methylated regions

Table S6 - List of IPA hits for genes corresponding to differentially bound regions and differentially methylated regions

Table S7 - Linear mixed model applied for estimation of lineage contribution as per each treatment group at 6 months and 12 months post-transplant.

Table S8 - Linear mixed model applied for estimation of human CD56+ satellite cells within the regenerating muscle for each treatment group at 1, 2, and 4 weeks post-transplant

Table S9 - Linear mixed model applied to Cohort A for estimation of host-derived (mouse) and chimeric muscle fibers for each treatment group at 4 weeks post-transplant.

Table S10 - Linear mixed model applied to Cohort B for estimation of host-derived (mouse), donor-derived (human) and chimeric muscle fibers for each treatment group at 8 weeks post-transplant.

### Supplementary methods

### Cell growth and self-renewal

#### *CFU-F assay*

Cells were seeded into 35 mm dishes and cultured in complete medium. The cells start forming colonies around day 8 in culture. At the end of 2 weeks, the cells in dishes were fixed in 4% paraformaldehyde (PFA) and washed twice in PBS for 15 minutes each. The fixed cells were then stained with Crystal violet (0.5% w/v in 80% ethanol) for 30 – 60 minutes, depending on the density of colonies. Dishes were then scanned at 600 dpi resolution and colonies quantified using Image J software [1].

#### *Long-term growth*

AdMSCs and iMS cells were expanded in bulk culture after plating 10,000 cells per T75 flask and passaged upon reaching 70-80% confluence. Expansion was done in either serum-free medium, FCS-supplemented medium or autologous/allogenic supplemented human serum as per the respective experiment. Cells were counted using a haemocytometer and cumulative cell numbers were calculated.

#### *Immunophenotyping*

The MSC phenotyping kit (Miltenyi Biotec) was used to evaluate cell surface marker expression. Briefly, adherent AdMSCs and iMS cells were harvested and labelled as per manufacturer's instructions and assessed by flow cytometry on a LSR Fortessa X20 cell analyser (BD Biosciences) with FacsDiva software (BD Biosciences). Gates were set using unstained and isotype stained controls and data was analysed using FlowJo software (Tree Star Inc., version 10.0.7).

### In vitro differentiation

The *in vitro* plasticity of control Ad-MSCs and reprogrammed iMS cells was determined by inducing the cells to undergo differentiation into various cell types using differentiation protocols adapted from a previous report [2]. Prior to induction, early passage (P3) AdMSCs and iMS cells were seeded at $2x10^4$ cells/well in a 6-well plate or $2x10^3$ cells/well in a 8-chambered slide and cultured in MSC medium until they reached confluence. During this period, medium was changed every 3-4 days.

#### *Osteogenic differentiation*

Well-adhered, confluent cells were switched to osteogenic medium containing αMEM (Life Technologies), 10% FCS, 100 μg/mL penicillin and 250 ng/mL streptomycin, 200 mM L-Glutamine, 0.1 μM dexamethasone (Sigma-Aldrich), 10 mM β-glycerophosphate. (Sigma-Aldrich), 200 μM L-ascorbic acid 2-phosphate (Sigma-Aldrich) for 21 days. The cells were then fixed in 4% PFA and stained for calcium deposition and extracellular matrix with freshly prepared 1% NH4+ buffered (pH 4.1-4.3) Alizarin Red S solution to determine osteogenesis.

#### *Adipogenic differentiation*

Well-adhered, confluent cells were switched to DMEM (Life Technologies, 11965092), containing 10% FCS, 100 μg/mL penicillin and 250 ng/mL streptomycin, 200 mM L-Glutamine and 0.5 mM methyl-3-isobutyl methylxantine (Sigma-Aldrich), 1 μM dexamethasone (Sigma-Aldrich), 6 μg/mL insulin (Sigma-Aldrich), 100 μM indomethacin (Sigma-Aldrich) for 7-10 days with medium changes every 3-4 days. Intracellular lipid droplets, which could be observed using light microscopy, were used to follow differentiation.

The cells were then fixed in 4% PFA and stained with Oil Red O that stains lipid droplets to determine adipogenesis.

#### *Chondrogenic differentiation*

Well-adhered, confluent cells were cultured in serum-free DMEM-HG, 100 µg/mL penicillin and 250 ng/mL streptomycin, 200 mM L-Glutamine, 50 µg/mL insulin-transferrin selenium (ITS) acid mix (BD Biosciences), 0.2 mM L-ascorbic acid 2-phosphate (Sigma-Aldrich), 1mM sodium pyruvate, 0.1 µM dexamethasone (Sigma-Aldrich), 40 µg/mL Proline (Sigma-Aldrich) and 10 ng/mL transforming growth factor β3 (TGF-β3; R&D Systems), and the medium was changed every 4 days. After 28 days in culture, differentiated cells were stained for sulfated proteoglycans with 1% Alcian blue to determine chondrogenesis.

#### *Endothelial differentiation*

Cells were cultured in MSC medium on chambered glass slides pre-coated with 0.1% gelatin. Once confluent, the cells were induced with DMEM-LG (Life Technologies, 11885084) supplemented with 2% FCS, 100 µg/mL penicillin, 250 ng/mL streptomycin, 200 mM L-Glutamine, 10ng/mL Vascular Endothelial Growth Factor (VEGF) and $10^{-8}$M dexamethasone, with medium changes every 3-4 days. For Matrigel assay, cells were plated on the chamber slides coated with Matrigel and cultured for 14 days. At the end of 14 days, tubes were fixed and stained in 4% PFA and stained for CD31, Platelet derived growth factor receptor β (PDGFRβ) and DAPI as detailed below.

#### *Myogenic (smooth muscle) differentiation*

Cells were cultured in MSC medium on chambered glass slides pre-coated with 0.1% gelatin. Once confluent, the cells were induced with DMEM-HG supplemented with 5% FCS, 100 µg/mL penicillin, 250 ng/mL streptomycin, 200 mM L-Glutamine and 50 ng/mL recombinant human platelet derived growth factor BB (rhPDGF-BB) (Miltenyi Biotec), with medium changes every 3-4 days. After 14 days, cells were fixed in 4% PFA and immunostained for smooth muscle myosin heavy chain (MYH1), alpha-smooth muscle actin (α-SMA) and nuclear stain DAPI as detailed below.

***Hepatocyte differentiation***

At 80% cell confluence, culture medium was switched to serum free DMEM-HG containing 100μg/mL penicillin, 250 ng/mL streptomycin, 200mM L-Glutamine, 20 ng/mL EGF (R&D Systems) and 10ng/mL of bFGF (R&D Systems) to inhibit cell proliferation for 2 days. After conditioning the cells, differentiation medium was added consisting of DMEM-HG supplemented with 20 ng/mL of HGF (R&D Systems) and 10 ng/mL of bFGF for 7 days. The cells were then cultured in DMEM-HG supplemented with 20ng/mL OSM, 1 μM dexamethasone, 10 μL/mL ITS premix and 100μg/mL penicillin and 250 ng/mL streptomycin for 14 days. Media was changed every 7 days. Hepatic differentiation was assessed by immunofluorescence staining for hepatocyte nuclear factor 4 alpha (HNF4α).

***Neuronal differentiation***

At 80% confluence, culture medium was switched to DMEM-HG medium containing 100μg/mL penicillin, 250 ng/mL streptomycin, 200mM L-glutamine and 1mM β-mercaptoethanol. Medium was changed every 3-4 days and cultured for 8-10 days. Neuronal differentiation was confirmed by expression of neuron specific TGFβ3.

*Immunocytochemistry*

Cells were grown on poly-l-lysine or gelatin coated 8 well MilliCell EZ slides and cultured in appropriate culture or differentiation medium. At the end of differentiation, cells were fixed in ice cold 4% PFA for 30 minutes, permeabilized with PBS containing 0.05% Tween-20 for an hour. The cells were washed with PBS and blocked in PBS containing donkey serum (v/v) for 1 hour at RT to minimise non-specific antibody binding. Cells were then washed and incubated overnight with primary antibodies diluted in PBS at 4°C in a humidified chamber. The following day, cells were washed in PBS and incubated in secondary antibody diluted in PBS for 1 hour at room temperature. Cells were washed again with PBS and stained with nuclear stain DAPI for 30 minutes. After a final wash with PBS, the slides were mounted with ProLong diamond antifade mounting reagent, which was allowed to cure for 24 hours prior to imaging. Fluorescence slides were imaged on a Zeiss LSM 780 confocal microscope using a 10x or 20x objective. Images were analysed and quantified using ImageJ. The antibodies used in these investigations are listed in Supplementary table S11.

**Animal Experiments**

***Labelling of AdMSCs and iMS cells***

Replication incompetent lentiviral vector LeGO iG2-Luc2 [3] was used for labelling cells with luciferase and green fluorescent protein (GFP). LeGo-iG2-Luc2 was co-transfected with packaging plasmids into HEK293T cells using Lipofectamine 2000 reagent (Life Technologies) and lentiviral supernatant was collected 48 hours after transfection, filtered and used for transduction of patient and passage-matched AdMSCs and iMS cells.

#### *Intervertebral disc injury model*

Lentiviral tagged 1x$10^6$ AdMSCs and iMS cells loaded onto Helistat collagen sponges were bilaterally implanted under anaesthesia into postero-lateral lumbar spine region bilaterally in immunodeficient NSG mice (n=9 for each group). All cell transplants were performed by a single orthopaedic surgeon. Animals were anesthetized using isoflurane (5% for induction, 2-3% for maintenance), then posterior midline incisions were made over the caudal portion of the lumbar spine and two separate fascial incisions were made 4 mm bilaterally from the midline. A blunt muscle splitting technique was used lateral to the facet joints to expose the transverse processes of L4 and L5 lumbar spines. The processes were then decorticated using a scalpel. Next, collagen sponges embedded with cells were implanted between the transverse processes bilaterally into the para-spinal muscle bed. Finally, the fasciae and skin were each closed using a simple continuous technique with Ligaclip Multiple Clip Applier (Ethicon) and betadine was applied at the suture site to prevent any infections. Post-surgery, the mice were monitored every day for a week and then three times a week until endpoint. In instances of delayed wound closure, the incisions were re-clipped. Animals were periodically imaged for bioluminescence to track the presence of transplanted cells. At the end of 3, 6 or 12 months, mice were sacrificed and their spines from the thoracic to caudal vertebral region were removed as a whole, including pelvis. The specimens were fixed in 4% PFA for 48 hours, decalcified in 14% EDTA, embedded in OCT and were cryosectioned sagittally at 5 μm thickness for histology and immunofluorescence analysis.

#### *Bioluminescence imaging*

Xenogen IVIS Spectrum CT system (Perkin Elmer) was used to capture bioluminescence. Briefly, 15 minutes after intraperitoneal injection of D-luciferin (150 mg/kg), images were acquired for 5 minutes and radiance (p/s/$cm^2$/r) was used for subsequent data analysis. For the intervertebral injury model, images were recorded every week for the first 3 months and then every alternate week until endpoint. For the teratoma experiment, images were recorded every week until study endpoint. The scanned images were analysed using the Living Image 5.0 software to evaluate changes in signal intensity which is a direct function of the number of luciferase- expressing cells present in the animal.

***Histology***

Cryosections were stained with hematoxylin and eosin to determine the overall tissue architecture and understand the scope of plasticity of the transplanted cells. Histology was also applied to evaluate presence of any ectopic tissue formation within the transplanted spines.

***Immunofluorescence***

Fixed tissue cryosections were permeabilized with PBS containing 0.05% Tween-20 for an hour, followed by blocking with 10% donkey serum/PBS before incubating with the respective primary antibodies ON at 4°C. The following day, samples were rinsed with PBS and then incubated with the secondary antibodies for 1 hr at 4°C. After washing three times with PBS, cells were incubated with DAPI for 20 minutes and then washed in PBS. The sections were briefly dried and then mounted using Prolong Diamond antifade mounting reagent.

***Muscle fibre counting***

Image analysis and muscle fibre counting was performed using ImageJ with the colocalization and cell counter plugins (NIH). Laminin staining was used to determine muscle fibre boundaries and Spectrin/hCD56 staining was used to determine contribution of donor cells to formation of regenerated muscle fibre.
